# Effects of larval density on gene regulation in *C. elegans* during routine L1 synchronization

**DOI:** 10.1101/284927

**Authors:** Io Long Chan, Oliver J. Rando, Colin C. Conine

## Abstract

Bleaching gravid *C. elegans* followed by a short period of starvation of the L1 larvae is a routine method performed by worm researchers for generating synchronous populations for experiments. During the process of investigating dietary effects on gene regulation in L1 stage worms by single-worm RNA-Seq, we found that the density of resuspended L1 larvae affects expression of many mRNAs. Specifically, a number of genes related to metabolism and signalling are highly expressed in worms arrested at low density, but are repressed at higher arrest densities. We generated a GFP reporter strain based on one of the most density-dependent genes in our dataset – *lips-15* – and confirmed that this reporter was expressed specifically in worms arrested at relatively low density. Finally, we show that conditioned media from high density L1 cultures was able to downregulate *lips-15* even in L1 animals arrested at low density, and experiments using *daf-22* mutant animals demonstrated that this effect is not mediated by the ascaroside family of signalling pheromones. Together, our data implicate a soluble signalling molecule in density sensing by L1 stage *C. elegans*, and provide guidance for design of experiments focused on early developmental gene regulation.

## INTRODUCTION

Developing *C. elegans* larvae can enter developmental arrest under unfavorable environmental conditions. For example, starvation-induced developmental arrest is reversible and can occur during all stages of larval development (Hu 2007; Seidel and Kimble 2011; BAUGH 2013; Schindler *et al.* 2014). While various developmental arrest paradigms serve as key models for studying a multitude of biological processes, *C. elegans* researchers also make use of such developmental arrest protocols to generate synchronous populations for experiments (Lewis and Fleming 1995). For example, rather than progressing through development, embryos that hatch as first larval stage (L1) worms in the absence of food will instead enter L1 developmental arrest. A routine method performed by *C. elegans* researchers makes use of this L1 arrest phenomenon to generate developmentally-synchronized populations for experiments. The general protocol is to bleach gravid adult animals to obtain embryos, followed by hatching and a short period of starvation in buffer to generate arrested L1s; arrested L1s can then resume development synchronously upon being provided with food (Emmons *et al.* 1979; Lewis and Fleming 1995).

However, details of the culture conditions used for population synchronization by L1 arrest – such as, e.g., worm density – are not commonly reported in the literature, and may have effects on organismal physiology. In the course of investigating dietary effects on gene regulation in *C. elegans*, we uncovered massive batch effects in an L1-stage RNA-Seq dataset. As the density of arrested L1 animals has been reported to affect starvation survival in an experimental setting that is similar to the routine population synchronization method (Artyukhin *et al.* 2013), we hypothesized that failure to control L1 arrest density in our initial studies resulted in the RNA-Seq batch effects.

We therefore set out to systematically characterize genome-wide effects of L1 density on gene expression using single worm RNA-seq. We characterized mRNA abundance genome-wide in two genetic backgrounds – wild-type N2 animals as well as the gonochoristic *fog-2* mutant – arrested in L1 at 1, 5, 20, or 100 eggs/μL. We identified 53 genes, primarily encoding various metabolic enzymes and potential signaling molecules, that were robustly affected by larval density in both strains. We confirmed these findings in detail for the relatively uncharacterized *lips-15* gene, generating a promoter *Plips-15::gfp* reporter strain as a sensor for L1 density. Finally, results from conditioned media experiments reveal that an ascaroside-independent pathway mediates this chemical communication between L1s, and suggest the possibility of an unknown density-sensing system in *C. elegans*. Together our results reveal a robust molecular phenotype that correlates with larval density during L1 developmental arrest, demonstrating that this variable must be carefully controlled when performing L1 arrest for molecular studies.

## RESULTS

### Effects of arrest density on gene expression during L1 starvation

To systematically identify density-regulated genes in arrested L1 larvae, we used low-input RNA-Seq (Ramskold *et al.* 2012; Trombetta *et al.* 2014) to characterize the transcriptome of individual L1 animals arrested at different densities (**Figure 1A, Supplemental Figure S1**). Briefly, we generated arrested L1s for experiments by bleaching gravid adults to collect embryos, and resuspended the asynchronous embryos at four different densities (1, 5, 20, 100 eggs/μL) in M9 media supplemented with polyethylene glycol 3350 (0.5%, w/v) to prevent animals from sticking to tips and tubes lacking food. Embryos hatched in M9 enter L1 arrest due to absence of food, and were collected as single L1s 22 hours later for RNA-seq (**Figure 1A, Methods**). As our initial studies were performed in the gonochoristic *fog-2* mutant background – in which hermaphrodites do not make sperm and are therefore phenotypically female (Schedl and Kimble 1988) – we repeated the dilution series and RNA-Seq in wild-type N2 animals to ensure that the results do not reflect physiological quirks of the *fog-2* mutant (**Supplemental Table S1, Supplemental Figure S2**).

**Figure 1.**
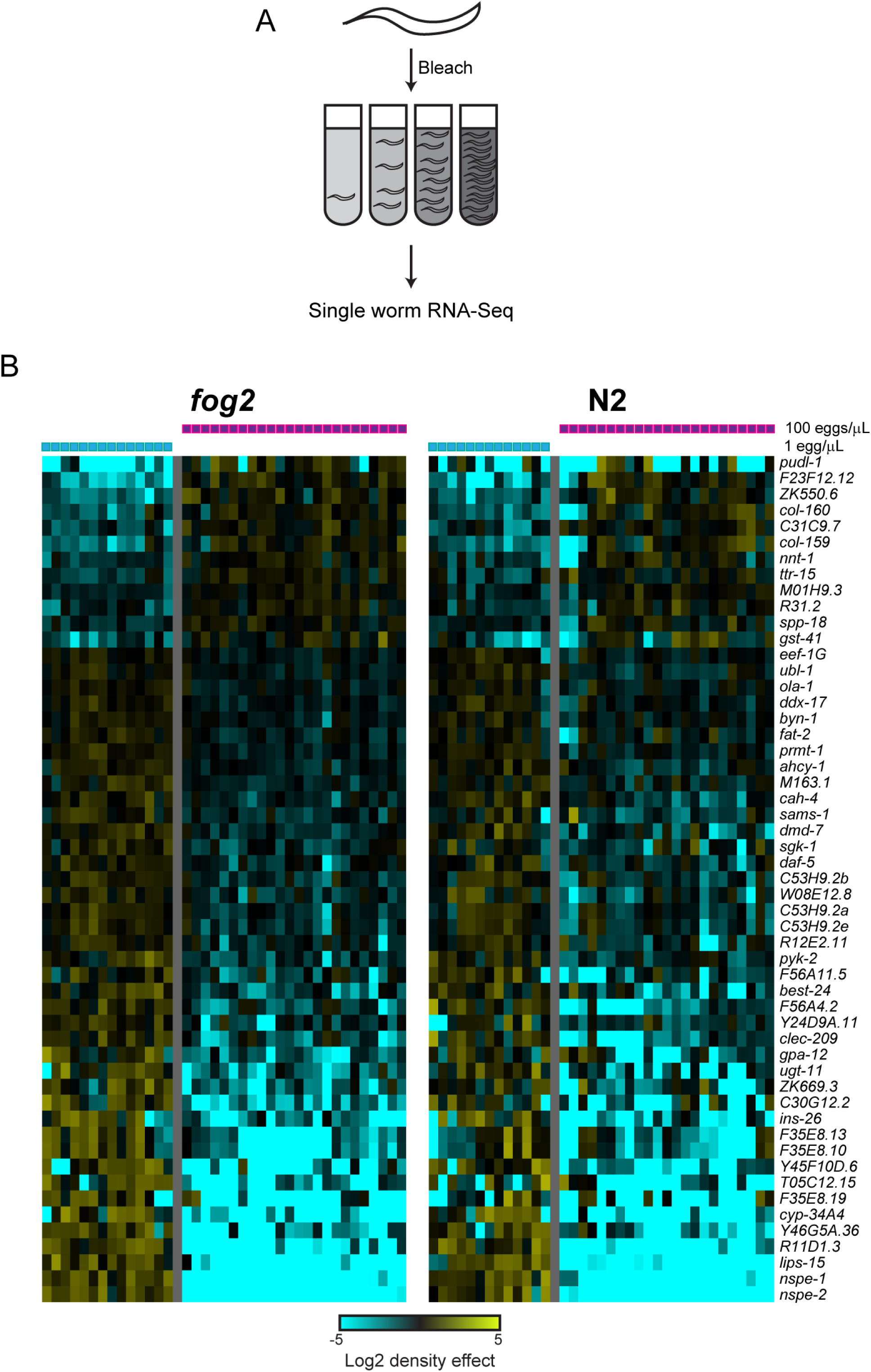
Effects of L1 arrest density on larval gene expression. A) Experimental schematic. Gravid *C. elegans* adults are bleached to recover a mixed population of early embryos. Embryos are then washed in M9 buffer containing PEG, counted, and resuspended at the indicated densities in 3 mL of M9. 22 hours post-bleaching, individual L1 animals were picked into separate tubes and processed for single-worm RNA-Seq. B) Significant effects of L1 arrest density on mRNA abundance. Heatmap shows clustered RNA abundance data for 53 genes significantly differentially-expressed (p_adj_ < 0.05) between 1 and 100 egg/μL cultures. Data are shown separately for replicate experiments using N2 hermaphrodites, or matings between *fog-2* mutant males and “females” (genotypically XX hermaphrodites which are incapable of making sperm and therefore phenotypically female), as indicated. In both datasets, mRNA abundance for individual L1s is expressed as the log_2_ ratio relative to the average for that gene across all individuals in the relevant strain background.

We first compared RNA-Seq profiles for L1s hatched at the highest and lowest densities (100/μL vs. 1/μL). As results in the *fog-2* and N2 backgrounds were essentially identical (**Supplemental Tables S2, S3**), we merged these datasets and identified a core set of 53 significantly density-regulated genes (**Figure 1B**). A small number of genes were upregulated under high-density growth arrest, including two collagen genes (many other collagen-related genes were also upregulated in these conditions but were not individually significant). In contrast to the moderate changes in gene expression for density-activated genes, several of the density-repressed genes were affected ∼10-100-fold by arrest density. Most notably, we find that four relatively uncharacterized genes located in a cluster on chromosome II – *nspe-1, nspe-2, lips-15*, and *lips-16*, encoding two predicted lipases and two potential signaling peptides – were highly-transcribed in low-density conditions, but were either undetectably expressed or expressed at low levels at high density. Other density-repressed genes encoded signaling molecules (*daf-5, ins-26, sgk-1, gpa-12, dmd-7*), metabolic enzymes (*sams-1, pyk-2, ahcy-1, fat-2, cah-4, cyp34-A4*), and a handful of uncharacterized genes (*F35E8.13*, etc.). Although we do not further characterize the physiology of animals maintained at high density, we predict that lipid metabolism – in particular, phosphatidylcholine metabolism (Walker *et al.* 2011) – is likely to be substantially altered in these animals.

### An L1 density reporter

To further validate the dramatic effects of arrest density on larval gene expression, we sought to develop a GFP reporter for L1 density. We focused on *lips-15*, which is highly-expressed under low density conditions (>1,000 tpm) and was nearly undetectable (median expression of 0 tpm, average of ∼20 tpm) in worms arrested at high densities. We therefore generated transgenic strains by integrating a promoter *Plips-15::GFP* fusion construct into the genome on chromosome II using MosSCI (Frokjaer-Jensen *et al.* 2008) (**Figure 2A**), again creating reporter lines in the N2 and *fog-2* backgrounds. We assessed the effects of L1 arrest density on reporter gene expression, resuspending reporter embryos at 1 or 100 eggs/μL. Consistent with the dramatic repression of *lips-15* mRNA abundance observed in high density L1 cultures, we confirmed that *Plips-15:*:GFP expression was strongly reduced in high density L1s (**Figures 2B-D**). Density-dependent regulation of this reporter was also observed in animals hatched into food, demonstrating that effects of density are not confined to conditions of starvation (**Supplemental Figure S3**).

**Figure 2.**
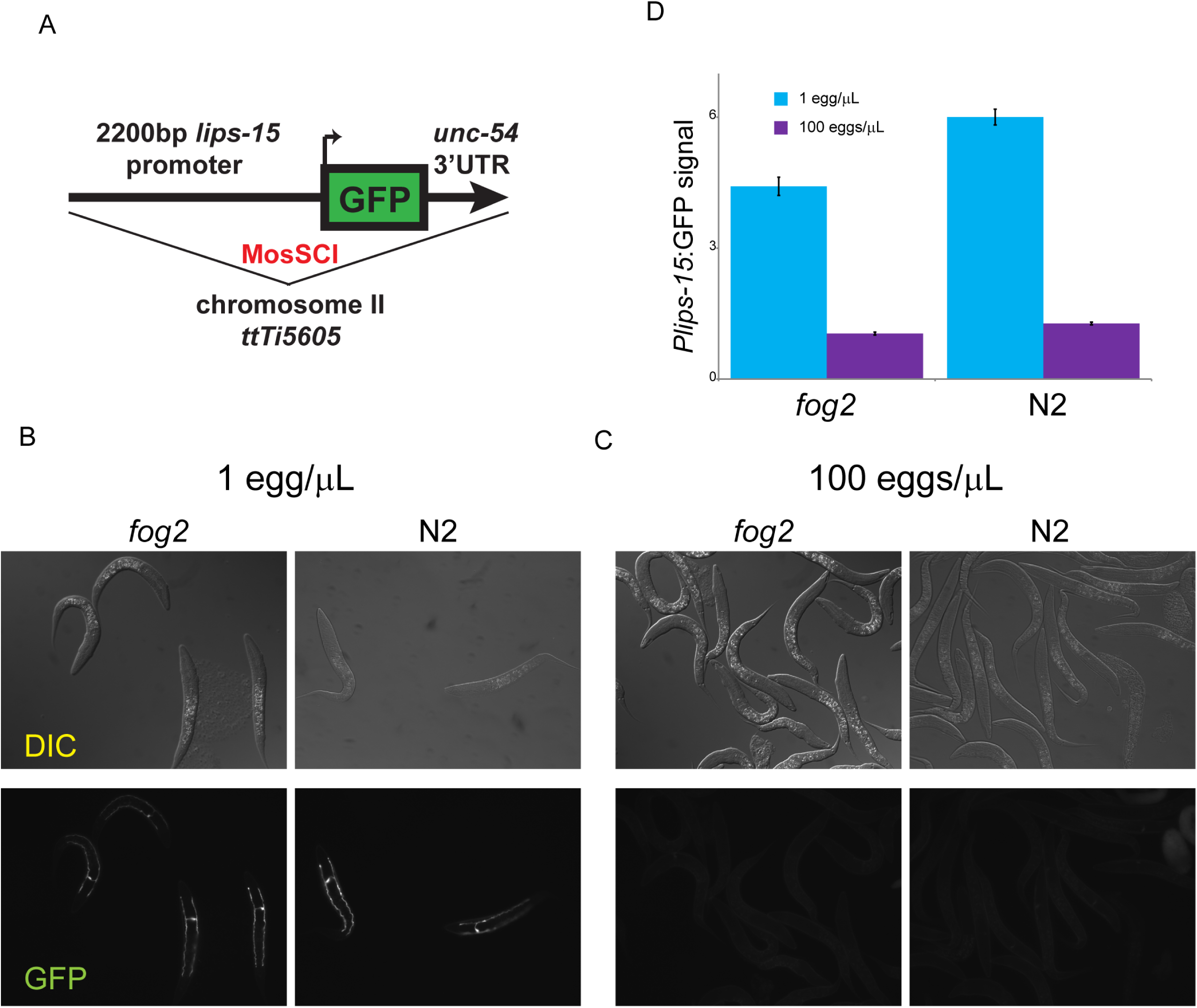
A GFP reporter for L1 density sensing. A) Schematic of the reporter for *lips-15* promoter activity. B) Typical images of *Plips-15::gfp* reporter expression in animals (of the indicated genetic background) arrested at low density. GFP expression is confined to a single large cell with long lateral projections, characteristic of the *C. elegans* excretory cell. C) As in (B), for reporter animals arrested at 100 eggs/μL. D) Quantitation of GFP expression at two densities in two strain backgrounds – data are shown as mean +/-s.e.m. for low and high density *fog-2* animals (n=206 and 230, respectively), and low and high density N2 animals (n=181 and 304, respectively).

Interestingly, the *Plips-15::GFP* reporter exhibits a surprisingly specific localization pattern at low density, with robust GFP expression confined to the excretory cell (**Figure 2B**). This cell serves a function analogous to the kidney of mammals (Nelson and Riddle 1984; Buechner *et al.* 1999), providing a further hint that the high density-repressed genes are involved in organismal metabolism. These results provide independent validation of our RNA-Seq dataset, and provide a robust single-worm reporter for L1 arrest density.

### Density signaling is mediated by a soluble factor

What is the nature of the signal that mediates density signaling? To further characterize the nature of the density-dependent signal, we examined the expression of density-regulated genes in animals arrested at intermediate L1 densities (5 and 20 eggs/μL). In general, we find that expression of these genes changes continuously across the four densities used, rather than exhibiting a switch-like transition at some density (**Figure 3**). Because our data are single-L1 RNA-Seq data rather than being an ensemble measure from pooled animals, the continuous changes in gene expression reveal a tuneable response being mounted in individual animals, rather than changes in the frequency of phenotypic subpopulations.

**Figure 3.**
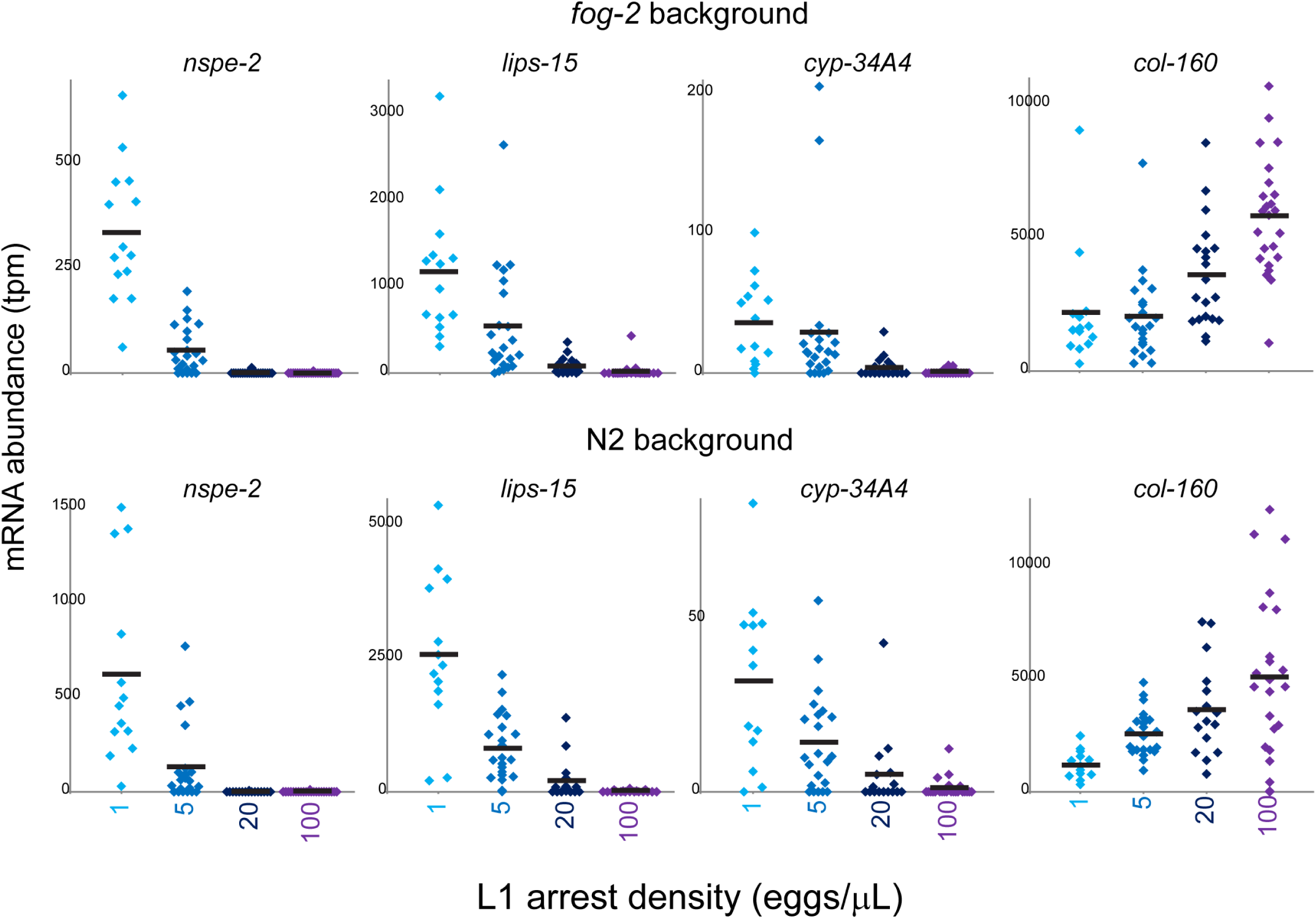
Continuous effects of arrest density on target gene expression. Dot plots show data for the indicated genes in individual L1s arrested at 4 different densities.

Is density signaling mediated by physical contact between animals, or by a soluble factor produced (or consumed) by worms? A previous study demonstrated that conditioned media generated from L1 larvae arrested at high density, provided to low-density L1s, is sufficient to phenocopy a long-term survival phenotype of L1s arrested at high density (Artyukhin *et al.* 2013). We therefore assayed *Plips-15::GFP* expression in animals bleached and maintained at low density in either control media or in conditioned media produced by L1-arrested animals cultured at high density (**Figure 4A**). As before, we observed robust expression of the reporter in animals maintained at low density in control media (**Figure 4B**). However, this expression was extinguished when L1s were arrested in high density conditioned media (**Figure 4C**). Repression by conditioned media was reversible, as 1) animals maintained at high density for 22 hours were capable of inducing *lips-15* expression when diluted 100X into control media conditions, and 2) animals hatched at low density in conditioned media were able to induce reporter expression after being washed and transferred to control media (**Figure 4D**). Activation of GFP occurred within hours in both cases, demonstrating that even under conditions of developmental arrest, L1 worms can sense and rapidly respond to changes in population density. Together with the observation that low density conditioned media could not activate *lips-15* in animals arrested at high density (not shown), our data reveal a density-sensing pathway that is mediated by a soluble factor produced by arrested *C. elegans* L1 larvae. We note that we cannot distinguish in this study between a signal produced only in animals maintained at high density, and a signal produced constitutively by L1 larvae but which is present in low density cultures at concentrations too low to activate the high density response.

**Figure 4.**
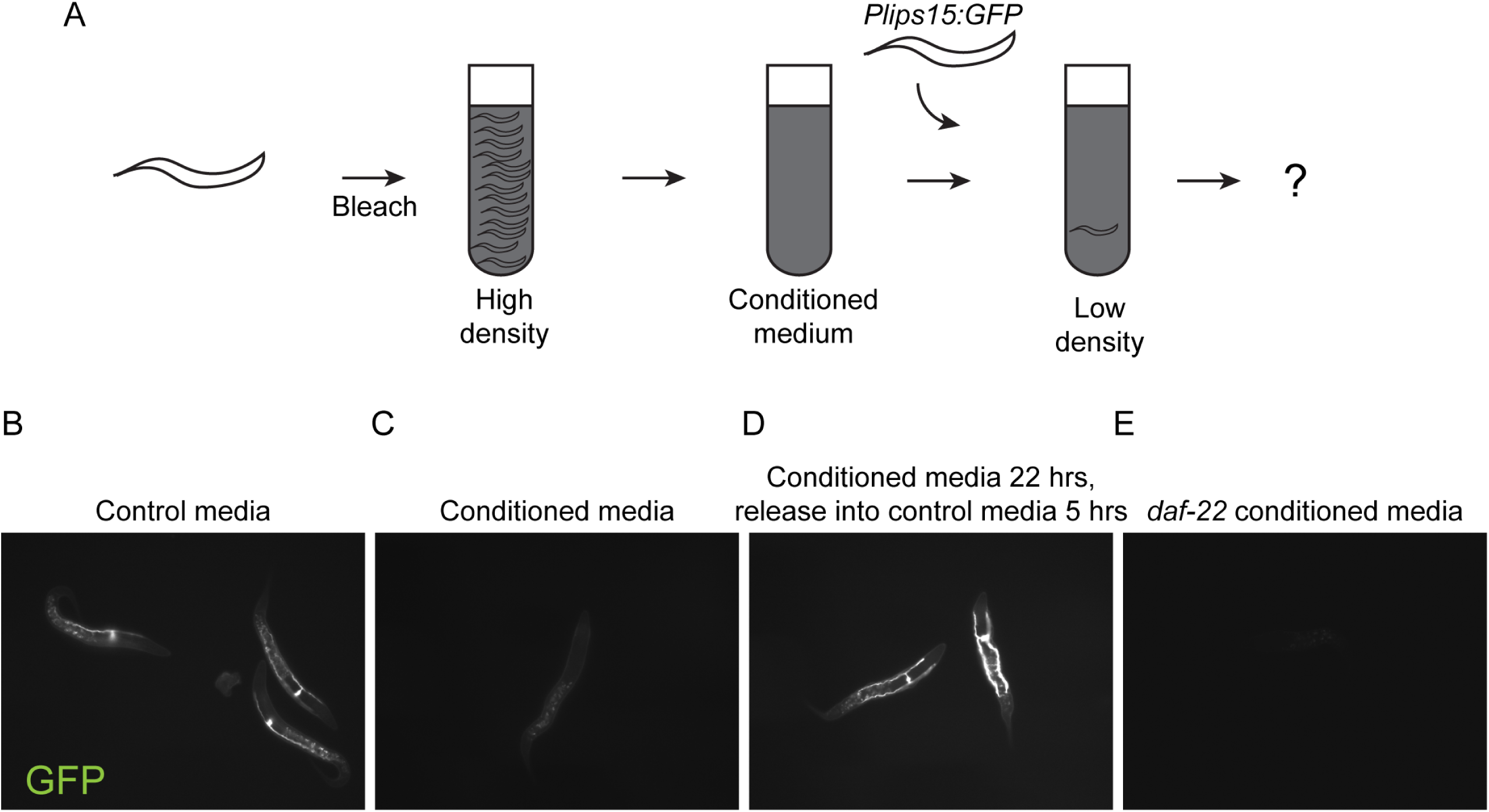
Density effects are mediated by a soluble, ascaroside-independent signal. A) Schematic of conditioned media experiments. B-E) GFP images for *Plips-15::gfp* animals maintained at 1 egg/μL in control media (B), high density L1-conditioned media (C), and conditioned media from high-density arrest cultures of *daf-22* mutant animals (E). In (D), reporter animals were arrested in conditioned media as in (C), then washed and suspended at low density in fresh control media for five hours.

### L1 density gene expression effect is mediated by a *daf-22*/ascaroside independent pathway

*C. elegans* development and behavior are regulated by a variety of small molecule signals. Prominent among these are the ascarosides, a diverse family of ascarylose sugar-based pheromones which regulate an array of phenotypes such as dauer entry and exit (Ludewig and Schroeder 2013). To test the hypothesis that the density effects observed here might be mediated by ascaroside signaling, we generated conditioned medium from high density L1 cultures of *daf-22* mutants, which do not produce any wild-type ascarosides (Butcher *et al.* 2009; Von Reuss *et al.* 2012). Conditioned media from *daf-22* mutants robustly repressed *lips-15* expression (**Figure 4E**), inconsistent with the hypothesis that ascarosides are responsible for this case of density signaling.

## DISCUSSION

Taken together, our data reveal that the density of starved *C. elegans* L1-stage larvae significantly affects gene expression, with some genes responding ∼10-100-fold to animal density. These data demonstrate that arrest density is an important experimental variable to control in molecular studies of L1-stage *C. elegans*. For example, recent pioneering studies of single-cell transcriptomes in developing *C. elegans* were characterized by substantial gene expression variability beyond the expected technical variation, which the authors suggested might be ascribed to developmental timing or preparation of the larvae (Cao *et al.* 2017).

### L1 density signaling in *C. elegans*

Several recent studies have shown that L1 arrest density can alter various phenotypes in *C. elegans*. For example, L1 density plays a role in parent-offspring signaling, as the presence of L1 larvae on plates was recently shown to alter food-related behaviors in adults (Scott *et al.* 2017). In this case, the relevant signal is lost in *daf-22* mutant, indicating that ascarosides are likely responsible for L1 density signaling. In contrast, an ascaroside-independent signal has been shown to mediate the effects of density on the survival of L1 worms under prolonged starvation (far longer than the time frame of our study) (Artyukhin *et al.* 2013). As in our study, the effects of L1 density on starvation survival could be transferred by conditioned media, and were not mediated by ascarosides. L1 arrest density has longer-lasting effects on worm physiology as well, as population density during larval development, but not density experienced as adults, alters later development rates and adult lifespan (Ludewig *et al.* 2017). As with density effects on L1 starvation survival, the density signal in this study was produced in *daf-22* mutants, again supporting the existence of a non-ascaroside signal produced by L1s.

It seems likely that the target genes identified here respond to this previously-described high-density signal, as we find that *lips-15* expression is suppressed by a density signal present in conditioned media from both wild-type and from *daf-22* mutants. Although we do not identify the signaling molecule(s) produced (or consumed) by L1 animals here, it is interesting to note that two of the most highly density-regulated genes – *nspe-1* and *nspe-2* – correspond to nematode-specific genes encoding predicted neuropeptides, and their response to animal density could reflect negative feedback control of their expression. Alternatively, the prominent regulation of genes related to lipid metabolism here could be related to the previously-reported antagonism between ascarosides and the high density signal (Ludewig *et al.* 2017), as for example it is plausible that secreted lipases (*lips-15*, etc.) or other enzymes could alter or degrade lipid signals. That said, the high density signal reported by Artyukhin *et al* appears to be a mixture of small (<3 kD) molecules, arguing against this latter hypothesis. Whatever the nature of the signal, the robust response of the *Plips-15::GFP* strain developed here could provide a convenient reporter for genetic screens to identify the relevant signal and its effector pathway.

## ACKNOWLEDGEMENTS

We thank L.R. Baugh, A. Artyukhin, M. Olmedo, and members of the Rando lab for helpful discussions. We thank members of the C. Mello and A.J.M. Walhout laboratory for sharing reagents, imaging equipment, protocols, and many helpful discussions. We thank C. Baer (UMass SCOPE facility), S. Guharajan, and Y. Shang for assistance with microscopy and image data analyses during the revision process. Some strains were provided by the CGC, which is funded by NIH Office of Research Infrastructure Programs (P40 OD010440). This work was supported by NIH R01 HD080224. CCC is a Merck Fellow of the Helen Hay Whitney Foundation.

## METHODS

### *C. elegans* strains and maintenance

All strains were maintained at 20 °C and passaged for at least four generations under non-starvation conditions before experimentation. The Bristol N2 strain was used as wild-type. Alleles used in this study listed by chromosome. LGII *daf-22(m130)* strain DR476, *Plips-15::gfp::unc-54 3’utr* (we are currently awaiting an allele and strain designation for these animals); LGV *fog-2(q71)* strain CB4108. The *Plips-15::gfp* reporter strain was constructed using MosSCI (Frokjaer-Jensen *et al.* 2008).

### Strain passaging and bleaching

Animals were passaged by bleaching gravid worms and directly plating 125,000 eggs mixed with *E. coli* OP50 (washed and resuspended in M9 buffer to 0.33 mg/mL), therefore worms never experience starvation under these conditions. Strains were maintained on 150 x 15 mm petri dishes that contained standard nematode growth media (NGM) with modification [10g agarose + 7g agar instead of 17g agar]. Gravid animals were washed from plates with M9 buffer, collected in conical tubes and centrifuged at 2000 x g for 30 seconds. Next, the supernatant was removed and worms were treated with 10 mL of bleach solution [41:6:3 ddH_2_O, sodium hypochlorite (Fisher Chemical, SS290-1), 5M KOH, and kept on a rocking platform. Worms were spun down after 4 minutes and the supernatant was replaced with fresh bleach. This procedure was repeated one more time and the suspension was visually checked to ensure all worm bodies were dissolved. The suspension was centrifuged at 2000 x g for 30 seconds, bleach solution was removed, and embryos were washed with M9 buffer supplemented with PEG 3350 (0.5%, w/v) three times before use. In this study, M9 buffer was always supplemented with PEG 3350 (0.5%, w/v) unless noted otherwise. The density of embryos in M9 were determined by manual counting.

For density experiments in the presence of food a single *E.coli* OP50 colony was picked and inoculated in 3 mL of LB media at 37°C for 4 hours, and this was used to inoculate 200 mL of LB media that was grown for 16 hours at 37°C. Next, the bacteria culture was pelleted at 3750 x g for 30 mins at 4°C and resuspended with M9, the final volume was adjusted to 10 mL including the appropriate number of embryos obtained by bleaching. Worms were cultured in 50 mL conical flasks at 20°C shaking at 180 rpm in a temperature-controlled incubator.

### Construction of the *Plips-15::GFP* L1 density sensor strain

The L1 density reporter strain was constructed using MosSCI with EG6699as the starting strain (Frokjaer-Jensen *et al.* 2008). The promoter region of *lips-15* was cloned using forward primer ATTTTTCACACAGAATTCCA and reverse primer with overhang into GFP GTGAAAAGTTCTTCTCCTTTACTCATGATGGAGTGAAGATTGTGGAG, and *GFP::unc-54 3’UTR* was cloned from genomic DNA of *Pacdh-1::GFP* (Arda *et al.* 2010) using a forward primer with homology to the promoter region of *lips-15* CTCCACAATCTTCACTCCATCATGAGTAAAGGAGAAGAACTTTTCAC, and reverse primer AAACAGTTATGTTTGGTATATTGG. The PCR products were gel purified and used as templates for PCR stitching. Next, the correct sized band was gel purified, inserted into a Zero Blunt TOPO cloning vector (ThermoFisher 450245), and fusion products from positive clones were determined by manual sequencing. The fusion product was then subcloned and inserted into the pCFJ350 plasmid, which was used for transformation by microinjection.

### Single L1 RNA-seq

Single L1 animals were placed in 5 ul of a typical worm lysis buffer (40mM Tris pH7.5, 10mM EDTA, 200mM NaCl, 0.5% SDS, and 0.4ug/ul Proteinase K), incubated at 55° for 10 minutes, and then frozen at −20°C until use for RNA-sequencing. For RNA-seq, a Smart-Seq2 protocol for generating cDNA followed by Nextera tagmentation was used to generate adapters and indexes required for sequencing (Trombetta *et al.* 2014). 96 samples (animals) were pooled and sequenced at a time on the NextSeq500 system (Illumina). Data were mapped using RSEM after removing rRNA and PCR duplicates.

For *fog-2* single L1 sequencing, 14 animals at 1 egg/μL passed our QC filters, with 22 animals at 5 eggs/μL, 20 animals at 20 eggs/μL, and 24 animals at 100 eggs/μL. For the N2 single L1 sequencing, animal numbers were 13, 22, 16, and 23. Average sequencing depth was 5.5 million reads/animal across all individuals, with ∼1 million reads typically aligning to the transcriptome (RSEM) after removal of PCR duplicates and rRNA reads (**Supplemental Table S1**).

### L1 conditioned medium experiment

High density conditioned medium was generated by first bleaching gravid animals grown on plates to obtain embryos, followed by resuspension of those embryos at high density (100/μL) in M9. At 24 hours, the actual density of hatched L1s was confirmed to be between 80-100/μL before processing for collection. To collect high density conditioned medium, we centrifuged the buffer containing L1s at high density at 2000 x g for 30 seconds, and passed the supernatant through a 1 μm glass fiber membrane syringe filter (Pall Life Sciences, Cat. # 4523T) to completely remove L1s and unhatched embryos. Filtered conditioned medium was stored at −80°C in 15 mL conical tubes (Corning #430791). Frozen Conditioned Medium was thawed in a 20°C water bath for at least 30 minutes prior to use.

To perform L1 conditioned medium experiments, embryos were obtained from *Plips-15::GFP* by bleaching gravid animals as described in this above. The density of embryos was counted and diluted to low density (1/ μL) in 3 mL of high density conditioned medium or control (M9) in 15 mL conical tubes. The tubes were placed on a rocking platform for 22 hours, and the density of L1s were confirmed before processing for imaging.

### Fluorescence imaging and quantification

*Plips-15::GFP* reporter animals were used for all imaging experiments. Animals were first hatched in the appropriate conditions, centrifuged at 2000 x g for 30 seconds then the pellet containing L1 worms was carefully removed and transferred to a PCR tube. Worms were treated with 1:10 volume of 10 mM tetramisole hydrochlorite (Sigma L9756) for 5 minutes before mounted onto a 2% agarose pad with a cover slip for Nomarski and fluorescence imaging using a Zeiss Axioplan2 Microscope.

For initial characterization of the reporter, animals were hatched in either low density (1/μL) or high density (100/μL) conditions in 3 mL of M9. For testing conditioned medium, animals were hatched at low density (1/μL) in either 3 mL of M9 (Control) or high density conditioned medium.

Image analysis was performed using Fiji/ImageJ to calculate background-corrected fluorescence, with the mean intensity of areas without worms or embryos taken as background, and the foreground signal calculated for the excretory cell bulb region.

## SUPPLEMENTAL MATERIALS

**Tables S1-S3.**

**Figures S1-S3.**

## SUPPLEMENTAL TABLES

**Table S1. Summary of sequencing from single L1 RNA-seq samples.** Mapping statistics for all individual worm RNA-Seq datasets in this study.

**Table S2. Fog-2 dataset.** Single-animal RNA-Seq data for *fog-2* animals arrested at 1, 5, 20, or 100 eggs/μL, as indicated.

**Table S3. N2 dataset.** Single-animal RNA-Seq data for N2 animals arrested at 1, 5, 20, or 100 eggs/μL, as indicated

## SUPPLEMENTAL FIGURES

**Figure S1.**
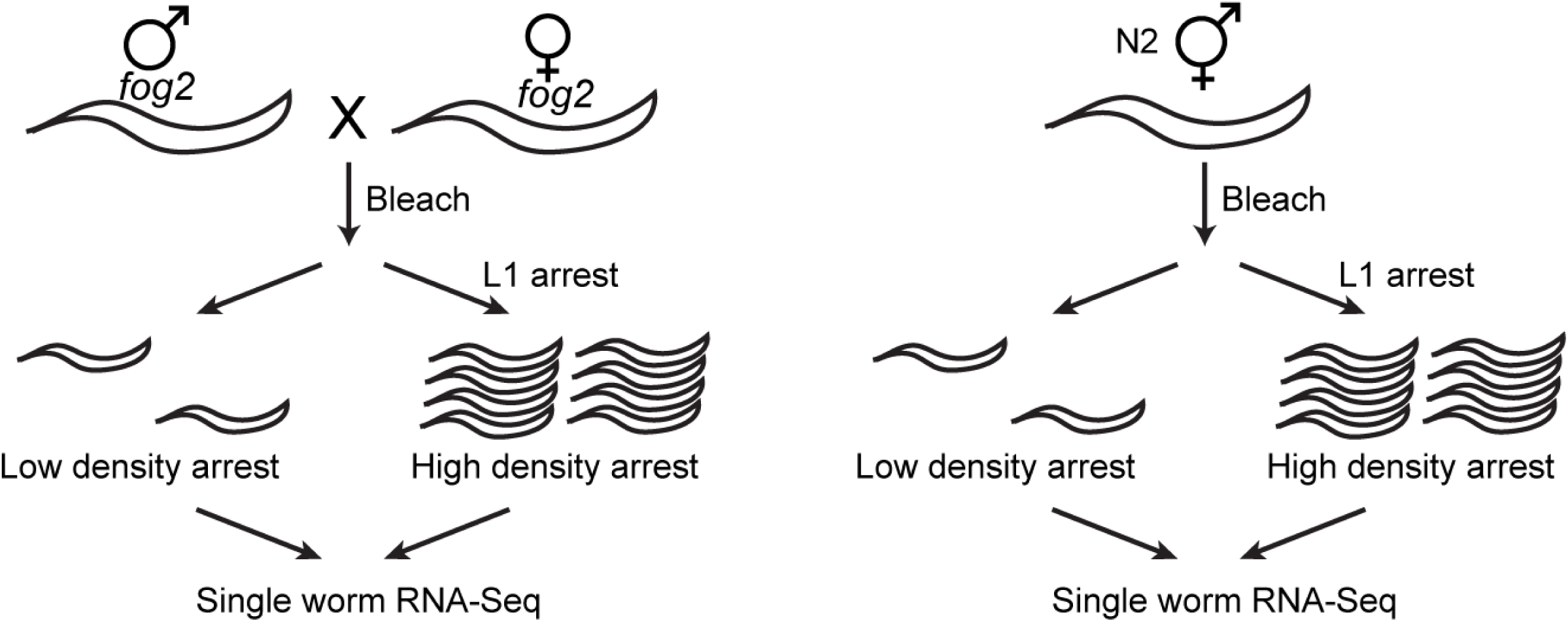
Experimental scheme. Comparison of N2 and *fog-2* background density experiments.

**Figure S2.**
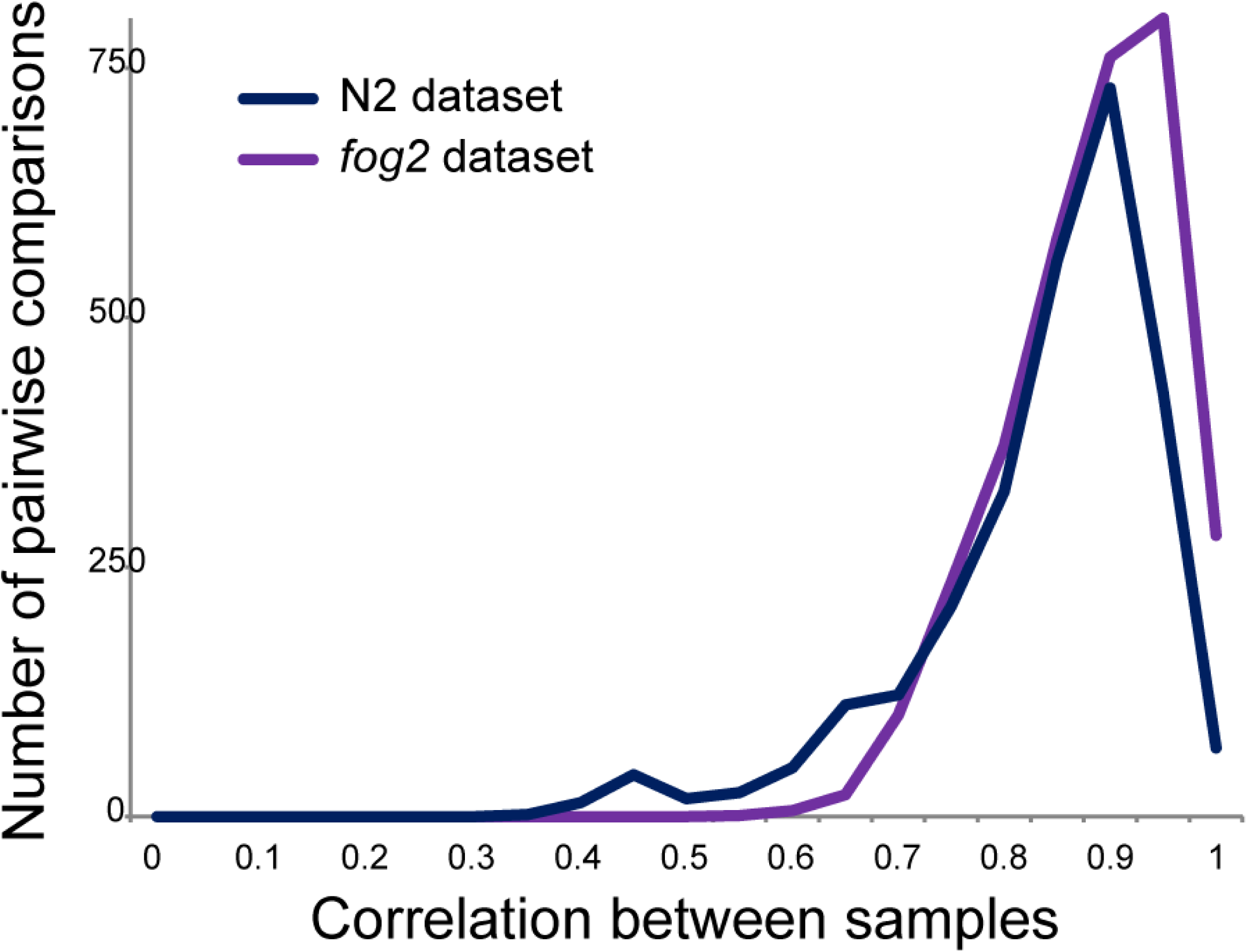
Deep sequencing reproducibility. For each experiment, we calculated pairwise correlation coefficients between individual sequencing libraries for all pairwise combinations of animals. Histograms show the distribution of correlation coefficients for each dataset, as indicated. Note the presence of a small number of outliers in the N2 dataset.

**Figure S3.**
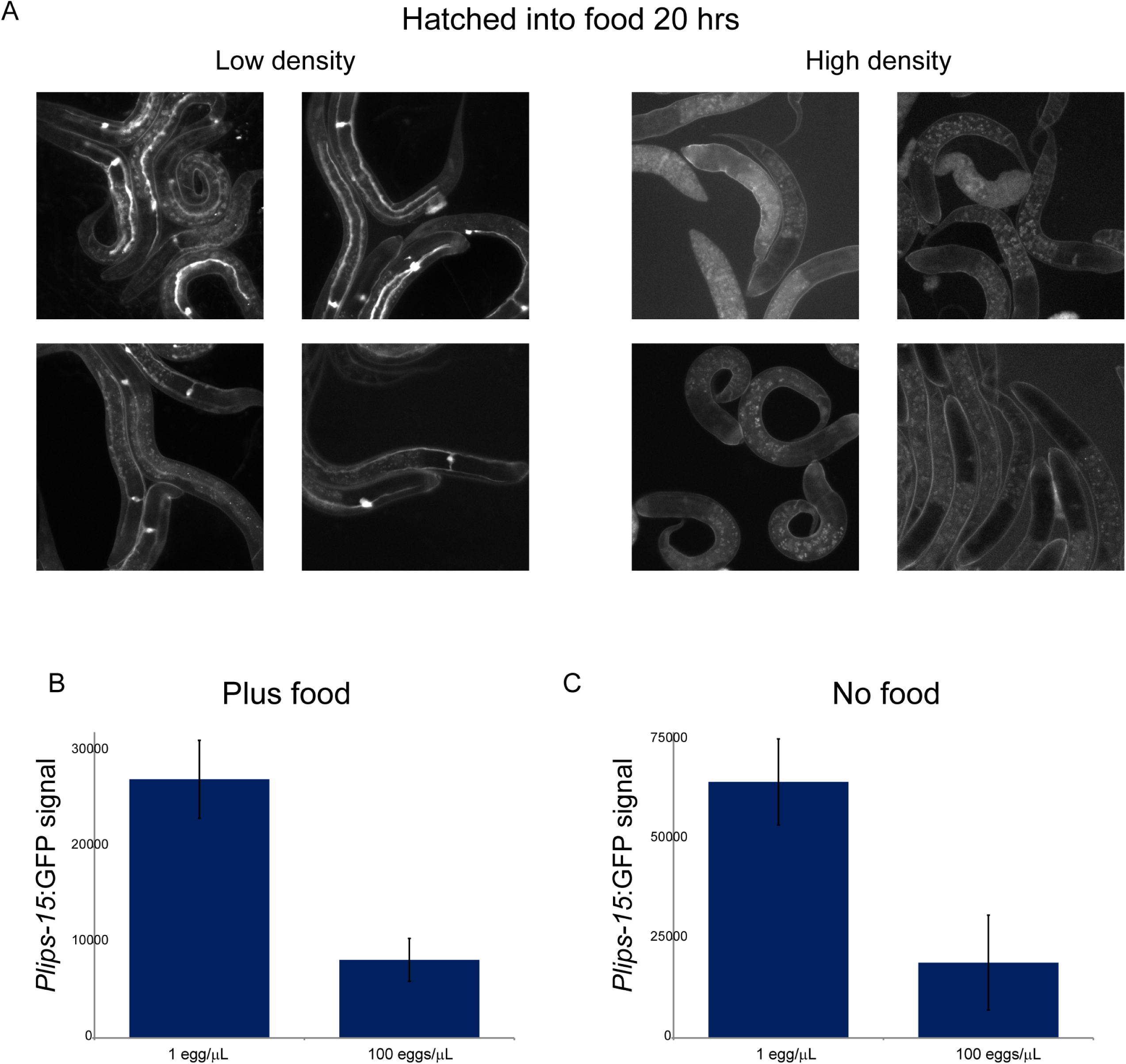
Density-dependent lips-15 repression is not starvation-dependent. A) GFP images for *Plips-15::gfp* animals at 20 hours post-bleaching at low density (1 egg/μL) or high density (100 eggs/μL) in M9 containing OP50 *E. coli* as food. B-C) Quantification of GFP expression at low (1 egg/μL) vs. high (100 eggs/μL) density, with food (B) or without food (C) as control. Data show mean +/-standard deviation of the background-corrected fluorescence intensity values for three biological replicate experiments, with 50-100 worms quantitated for each biological replicate.

